# Structural guidance in 3D microgel-in-hydrogel systems to improve chondrogenesis

**DOI:** 10.64898/2025.12.19.694877

**Authors:** Julia Kamp, Alisa C. Suturin, Claudia A. Garrido, Xiaole Tong, Jietao Xu, Nicole Kops, Lea-Sophie Müller, Till Hülsmann, Laura Klasen, Tamás Haraszti, Gerjo J.V.M. van Osch, Laura De Laporte

## Abstract

Anisotropic biomaterials are broadly studied in regenerative medicine as they aim to mimic the hierarchical structures of native tissue. The Anisogel, an anisotropic microgel-in-hydrogel system, involves two key factors for tissue engineering: guidance cues – magnetically aligned rod-shape structures - and a surrounding matrix - polyethylene glycol-vinylsulfone (PEG-VS) with a degradable peptide crosslinker. This research shows the potential of using a PEG-VS-based Anisogel to allow for guided ingrowth and chondrogenic differentiation of human mesenchymal stromal cells (hMSCs) modulated by the alignment and size of anisometric microgels. The properties of the surrounding matrix are optimized for the availability of anchoring peptides (RGD concentration) and matrix density (hydrogel precursor concentration). The microgels of highest tested dimensions (10×10×100 µm^3^) lead to a donor-independent significant upregulation of chondrogenic markers (*SOX 9*, *ACAN* and *COL2AI*) and a decrease of hypertrophic chondrogenic markers (*RUNX 2*), compared to unaligned and smaller microgel sizes. As a proof of concept, the Anisogel is tested in a semi-orthotopic mouse model where the effect of microgel alignment on cell infiltration and osteochondral tissue formation is evaluated.

## 1. Introduction

Articular cartilage is a highly specialized tissue consisting of distinct zones differentiated by tissue organization and extracellular matrix (ECM) composition. The tissue can be subdivided into the superficial, transitional, and deep zones, with each zone playing a critical role in the mechanical and biological functionality of the tissue.^[1,2]^ The complex structure, low cellularity in the deeper zones and the absence of blood vessels in articular cartilage limit its self-repair capabilities and represent a challenge for tissue regeneration.^[3–5]^

To promote cartilage regeneration, hydrogels are often employed to provide mechanical support for cells and introduce specific biochemical and physical cues to guide cell fate and behavior.^[6]^ Natural hydrogels based on either proteins or polysaccharides such as gelatin, hyaluronic acid, or chondroitin sulfate (CS) are employed in cartilage engineering due to their good biocompatibility and biodegradability.^[7–10]^ To improve the ease of tunability in physical and biochemical cues and decrease the risk of immunogenicity, synthetic hydrogels such as polyethylene glycol (PEG) have been increasingly used. It was previously shown that mesenchymal stromal cells (MSCs) can undergo chondrogenesis and produce cartilaginous tissue when embedded into PEG hydrogels.^[11–14]^ Furthermore, the impact of varying the concentration of binding motifs and the stiffness of these biomaterial platforms on chondrogenic differentiation have been depicted.^[15]^ While significant advances to regenerate cartilage tissue have been made over the last decades, most of these approaches aim at homogenous tissue formation mimicking the overall bulk properties and thereby not resembling the native-like structure.

Due to the functional importance of the hierarchical structure of cartilage, several approaches have been published that aim to mimic this organization by using multi-layered hydrogels. Reconstructing the layered structure of articular cartilage has been achieved by optimizing spatially patterned materials^[16]^, stiffness gradients^[17, 18]^ or by improving bioprinting strategies^[19]^. Beyond adjusting the material properties itself, other working groups have explored the use of different cell types for zonal cartilage repair by using chondrocytes harvested from specific zones^[20]^, or co-cultures of chondrocytes and osteogenic progenitors^[21]^. Guiding structures, such as collagen fibrils^[22]^ or microstructures (e.g., microribbons)^[23]^, are crucial for cell guidance and alignment, and have shown to be cell instructive structures that induce maturation of cartilaginous tissue.

Despite these advances in obtaining new cartilage tissue resembling the zonal-specific compositions, these constructs still lack the native tissue organization. To achieve ECM fiber alignment, early methods to organize ECM fibers relied on a strong magnetic field to align, for example, collagen fibers (2-10 T).^[24, 25]^ Over the past few decades, a broader toolbox has been developed, including flow-induced fiber alignment, stretching gels to induce fiber organization, or embedding magnetic particles that restructure gels by traveling toward an external magnet.^[26]^ However, these methods are not compatible with many hydrogels, usually require the implantation of pre-aligned constructs, and rely on high magnetic fields or high concentrations of potentially cytotoxic magnetic particles. Thus, there is a pressing need for new, modular, anisotropic hydrogel systems that can be crosslinked *in situ* without relying on strong external fields.

To fill this gap, our group has established a microgel-in-hydrogel system, called the Anisogel, that can be combined with any hydrogel material. It can be crosslinked *in situ* and only requires a weak magnetic field during crosslinking to induce anisotropy.^[27]^ The Anisogel is a structured hydrogel that allows for directed cell growth and controlled, hierarchical ECM protein deposition in a 3D environment.^28^ Magneto-responsive, rod-shaped guiding elements (PEG microgels or short polymeric fibers embedded with small amounts of superparamagnetic iron oxide nanoparticles (SPIONs)^[29]^) are suspended in the hydrogel precursor solution. By the application of a low magnetic field (< 100 mT), the rod-shaped guiding elements align with the magnetic field lines. Upon hydrogel gelation, the microgel or fiber orientation is fixed and the magnetic field can be removed. It has been shown that this material guides unidirectional cell growth when using fibroblasts^[28, 30, 31]^, vascular cells^[32]^, and nerve cells^[33–36]^.

This study used the Anisogel to guide cartilage regeneration by mimicking the structure of native tissue. To study the effect on chondrogenesis, human bone-marrow derived MSCs were encapsulated in an enzymatically degradable PEG hydrogel, modified with different peptides and embedded with different microgels. The surrounding hydrogel is made from 4-armed PEG-vinylsulfone (PEG-VS) that is crosslinked via a thiol-Michael type addition reaction with a matrix metalloproteinase (MMP)-sensitive peptide crosslinker, containing two thiol-bearing cysteines. The surrounding hydrogel matrix is developed to support cell spreading and chondrogenesis by optimizing the concentration of the cell-adhesive peptide GRGDSPC and the hydrogel stiffness. In a next step, magnetic, rod-shaped PEG microgels are incorporated to evaluate the effect of microgel alignment and size on cell alignment and chondrogenic differentiation. Transcriptional upregulation of chondrogenic markers - SRY-Box Transcription Factor 9 (*SOX9*), aggrecan (*ACAN*), and collagen II (*COL2A1*) - was most pronounced with the largest tested microgel dimensions (10×10×100 µm³) when aligned, showing a donor-independent impact of these guiding elements on early hMSCs chondrogenesis. The optimized microgel-in-hydrogel system was then evaluated *in vivo* using a semi-orthotopic mouse model. Here, an osteochondral defect was made in a biopsy of bovine articular cartilage with underlying bone and filled with the microgel-in-hydrogel system before being implanted subcutaneously in athymic mice.^[37, 38]^ New tissue formation was investigated for different microgel orientations (horizontal, vertical, and random). Our research shows the potency of anisotropic materials in guiding hMSCs chondrogenic differentiation, which can further contribute to regenerative medicine research.

## 2. Results and Discussion

### 2.1 The surrounding hydrogel matrix

The hydrogel precursor solution consists of a 4-armed PEG-VS (20kDa) and is crosslinked via Michael-type addition with a short, enzymatically MMP degradable peptide (GCRE**GPQG`↓` IWGQ**ERCG-NH_2_, cleavage site indicated by **`↓`**).^[39, 40]^ The resulted matrix can be produced with variable stiffness, is degradable on cell-demand, and contains reactive binding sites for biomolecules. In previous studies done by others, it was demonstrated that an RGD concentration of 150 µm enhanced the spreading and GAG content of human periosteum derived cells (hPDCs) when compared to PEG-VS hydrogels without RGD independent of the hydrogel stiffness.^[15]^ Therefore, we first optimized this hydrogel for human bone marrow-derived MSCs (hMSCs) in 3D cell culture by varying the RGD and polymer concentration. First, RGD concentrations were varied from 150 µM to 2 mM in a 6.5 wt% hydrogel (**Figure 1**) and hMSCs were incorporated into the hydrogel at 2 million cells mL^-1^ and cultured for 3 weeks in either control medium with 10% FBS or in serum-free chondrogenic differentiation medium with dexamethasone (10^-7^ M) and TGF-ß (10 ng/mL) (**Figure 1 B**). The highest 2 mM RGD concentration led to the most cell spreading, displayed by a higher projected cell area (**Figure 1 C**) and lower circularity (**Figure 1 D**). Chondrogenic differentiation was obvious from aggrecan and collagen II immunohistochemistry (**Figure 1 B**). Cells had a higher deposition of aggrecan and collagen II when encapsulated in hydrogels with higher RGD content (2 mM) compared to a lower RGD content (**Figure 1 E-F**). Interestingly, the differences in aggrecan and collagen II deposition between the control and differentiation media were not statistically significant. This suggest that the material properties may already induce chondrogenesis.

**Figure 1.**
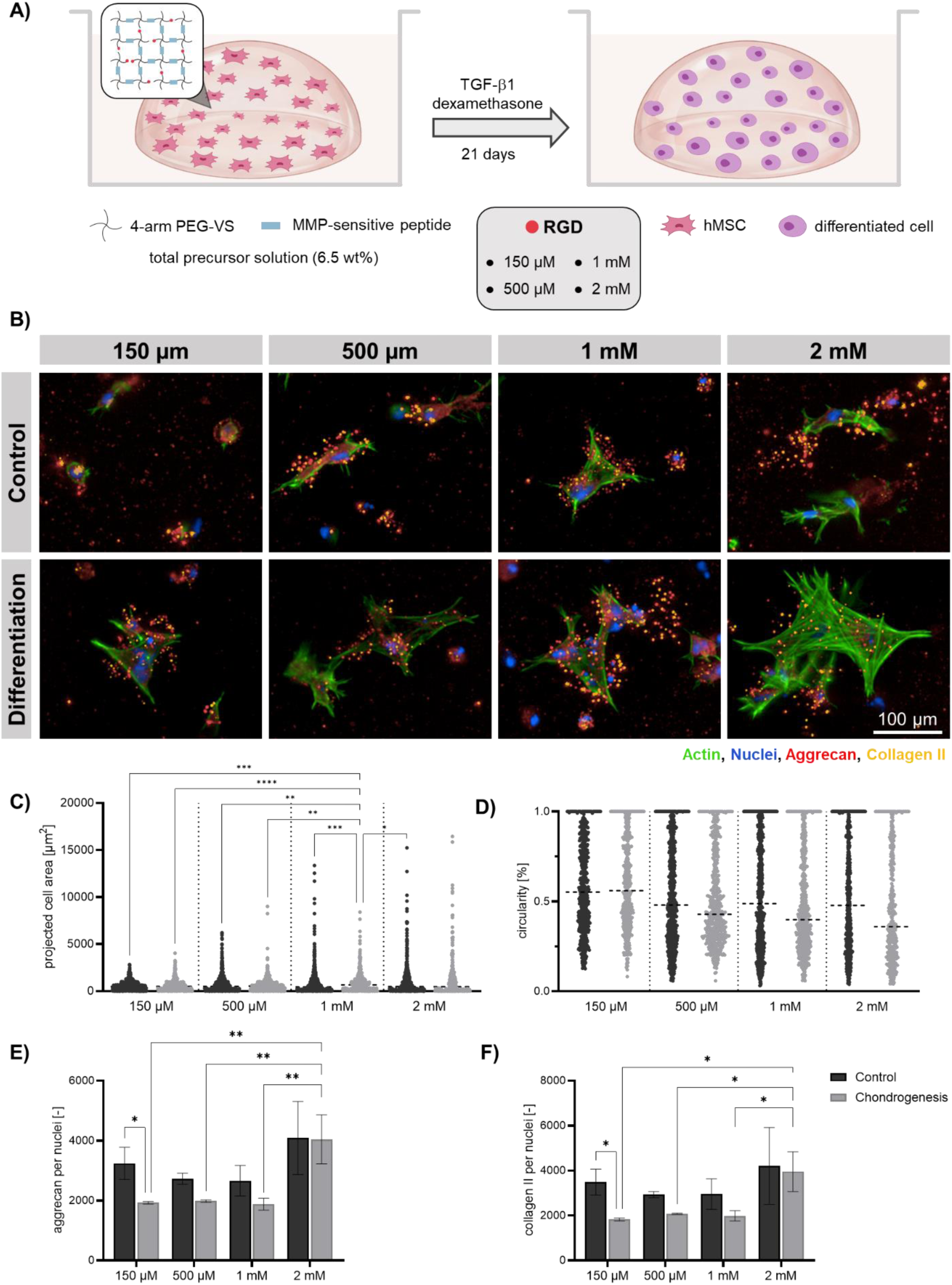
Optimization of RGD concentration in PEG-VS hydrogels (6.5 wt%) for chondrogenesis. A) Schematic protocol of hMSC differentiation in PEG-VS hydrogel droplets with varying RGD concentration. B) Representative confocal microscopy images of hMSCs stained for F-actin (green), DAPI (blue), aggrecan (red) and collagen 2 (yellow) cultured in control and differentiation conditions in hydrogels containing 150 µM, 500 µM, 1 mM, or 2 mM RGD. C,D) Cell spreading quantification in 10X maximum projection images of a 500 µm z-stack, cell area (C) and circularity (D). E,F) Quantification of confocal images for chondrogenic protein production for aggrecan (E), and collagen II (F). Data are shown as mean ± SD from three independent measurements (n = 3). Statistical significance was determined using one-way ANOVA followed by Tukey’s multiple comparisons test. Significance is indicated as: ns (p ≥ 0.05), * (p < 0.05), ** (p < 0.01).

To find the optimal hydrogel physico-mechanical properties, we used 3.4 wt%, 5 wt%, 6.5 wt% or 10 wt% PEG-VS hydrogels with 2 mM RGD (**Figure 2**). All hydrogels were produced under stochiometric conditions between the number of reactive vinylsulfone (from PEG-VS) and thiol groups (from MMP-sensitive peptide). The hydrogel weight percentage has an impact on the stiffness, porosity, and degradation rate of the hydrogel. Rheological characterization of the hydrogels measured storage moduli ranging from 0.5 kPa to 5 kPa, depending on the polymer content (**Figure 2 C**). After 3 weeks in culture, the production of the chondrogenic proteins aggrecan and collagen II was investigated via immunohistochemical analysis (**Figure 2 B**). Both proteins, aggrecan (**Figure 2 D**) and collagen II (**Figure 2 E**), were significantly higher for the softest 3.4 wt% PEG hydrogels, with a slight increase in the case of differentiation media. Softer materials (with a lower percentage of PEG-VS) were not considered as they are degraded too fast and do not last the complete culture time (3 weeks).

**Figure 2.**
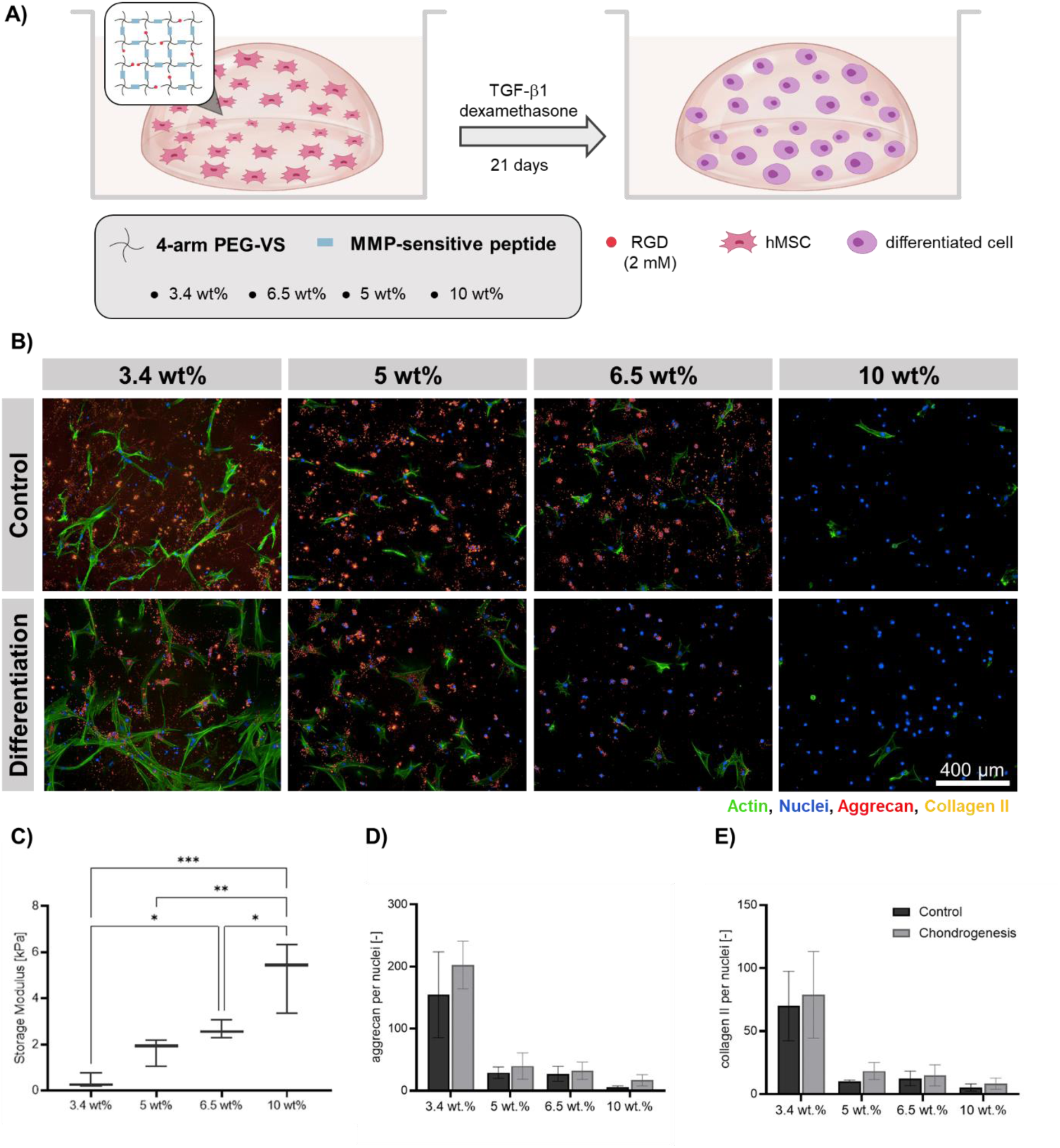
Optimization of hydrogel concentration for PEG-VS hydrogels with 2 mM RGD. A) Schematic protocol of hMSC differentiation in PEG-VS hydrogel droplets with varying precursor concentration. B) Confocal microscopy images of hMSCs stained for F-actin (green), DAPI (blue), aggrecan (red) and collagen 2 (yellow), and cultured in control compared to differentiation conditions in 3.4 wt%, 5 wt%, 6.5 wt%, or 10 wt% PEG-VS hydrogels. C) Storage moduli obtained via rheological measurements. Data are shown as mean ± SD from three independent measurements (n = 3). Statistical significance was determined using one-way ANOVA followed by Tukey’s multiple comparisons test. Significance is indicated as: ns (*p* ≥ 0.05), * (*p* < 0.05), ** (*p* < 0.01), *** (*p* < 0.001). E), F) Quantification of confocal images using 10X maximum projection images of a 500 µm z-stack for chondrogenic protein production for aggrecan (E), and collagen II (F).

These results are consistent with literature showing that matrices, produced by other materials, with a stiffness around 0.5 kPa are optimal for hMSCs chondrogenesis.^[41–43]^ Interestingly, in the case of different cell types, like human periosteum-derived cells (hPDCs) and ATDC5 cells, stiffer 6.5 wt% PEG-VS hydrogels demonstrated the highest level of chondrogenesis.^[15]^ This could be related to the initial cell loading or by the MMPs produced by the cells incorporated.

### 2.2 Implementation of magnetic rod-shaped microgels to guide chondrogenic differentiation

In the next step, magnetic rod-shape microgels of different dimensions were mixed and aligned inside the optimized PEG-VS hydrogel (3.4 wt% PEG-VS, 2 mM RGD) to form Anisogels. We explored different microgel sizes (2.5×2.5×25, 5×5×50, 10×10×100 µm^3^) to test their effect on hMSCs growth, alignment, and chondrogenic differentiation (**Figure 3**). The microgels were produced by in-mold polymerization as previously reported using PEG-diacrylate, and are bioinert as they are not modified with any bioactive molecule. They were simple embedded in the precursor solution at a volume percent of 1%, together with the cells at a concentration of 2 million cells per mL^-1^, aligned by the placement of magnets outside the droplets, and fixed in place by crosslinking of the surrounding hydrogels. The relation between the microgels’ dimensions, amount, volume concentration in the precursor solution, and inter-microgel distance can be found in a previous report.^44^ The interaction between hMSCs and rhodamine labelled microgels was depicted using confocal microscopy (**Figure 3 B**). The images reveal that alignment of microgels changed the cell morphology and spreading. The evaluation of the cell alignment showed that cells aligned preferentially along the microgel direction, independent of the microgel size, while unaligned microgels (**Figure 3 C**) resulted in random cell orientation precursor solution before gelation. The effect of cell alignment seems to be enhanced by the addition of biochemical signals (e.g., TGF-β in the chondrogenic media), as cells in control samples showed less collective orientation (**Figure S1**). It is important to highlight that the morphology of the encapsulated hMSCs did not resemble the morphology of native chondrocytes, as the *in vivo* chondrocyte morphology is dependent on the strain they experience during physiological mechanical loading of the cartilage^[45]^, and the results shown in this research are based on 3D culture systems that were not subjected to any type of external mechanical stimulation.

**Figure 3.**
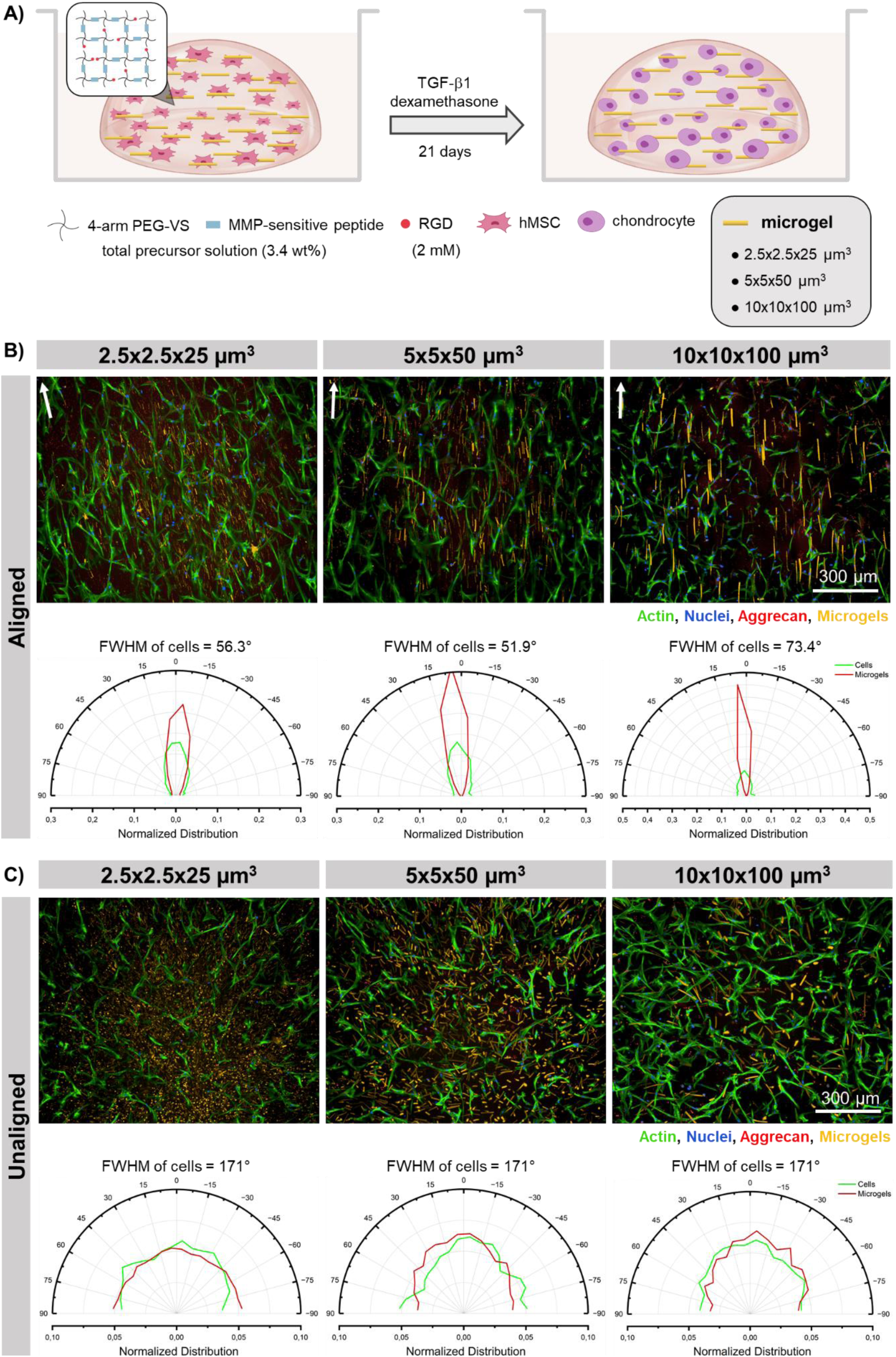
Varying the microgel dimensions and alignment of PRINT microgels in optimized PEG-VS hydrogel. A) Schematic protocol of hMSC differentiation in optimized PEG-VS hydrogel droplets for microgels with dimensions of 2.5×2.5×25 µm^3^, 5×5×50 µm^3^, or 10×10×100 µm^3^. Representative confocal microscopy images of hMSCs in hydrogels stained with F-actin (green) and DAPI (blue) cultured in optimized PEG-VS hydrogels in chondrogenic media used for the correspondent angle histograms depicting the full width at half maximum (FWHM, of n=1 hydrogel containing n>80 cells in the field of view) of cell (green) and microgel (red), in B) aligned and C) unaligned microgels.

Besides the cell morphology and alignment, the microgel size and orientation influenced chondrogenic differentiation (**Figure 4**). The expression of the chondrogenic markers aggrecan (*ACAN*, **Figure 4 A**), collagen II (*COL2A1*, **Figure 4 B**), and SRY-Box Transcription Factor 9 (*SOX9*, **Figure 4 C**) were evaluated using quantitative polymerase chain reaction (qPCR). The aligned microgels led to significantly higher expression of *SOX9*, *ACAN* and *COL2A1* (**Figure 4 A-C)**, mainly for the larger microgels, for the smallest microgels alignment did not influence *ACAN* and *COL2A1*. This is the first report demonstrating the impact of aligned short micro-guiding structures on chondrogenesis. The low expression of *COL2A1*, whereas the high expression of *SOX9* and *ACAN* reveal the early stages of chondrogenic differentiation. These two markers are expressed during the aggregation and early condensation, where, MSCs initially cluster through cell-cell adhesion (aggregation) and then become densely packed, facilitating the cellular signaling and extracellular matrix changes necessary for cartilage formation (condensation).^[46]^

**Figure 4.**
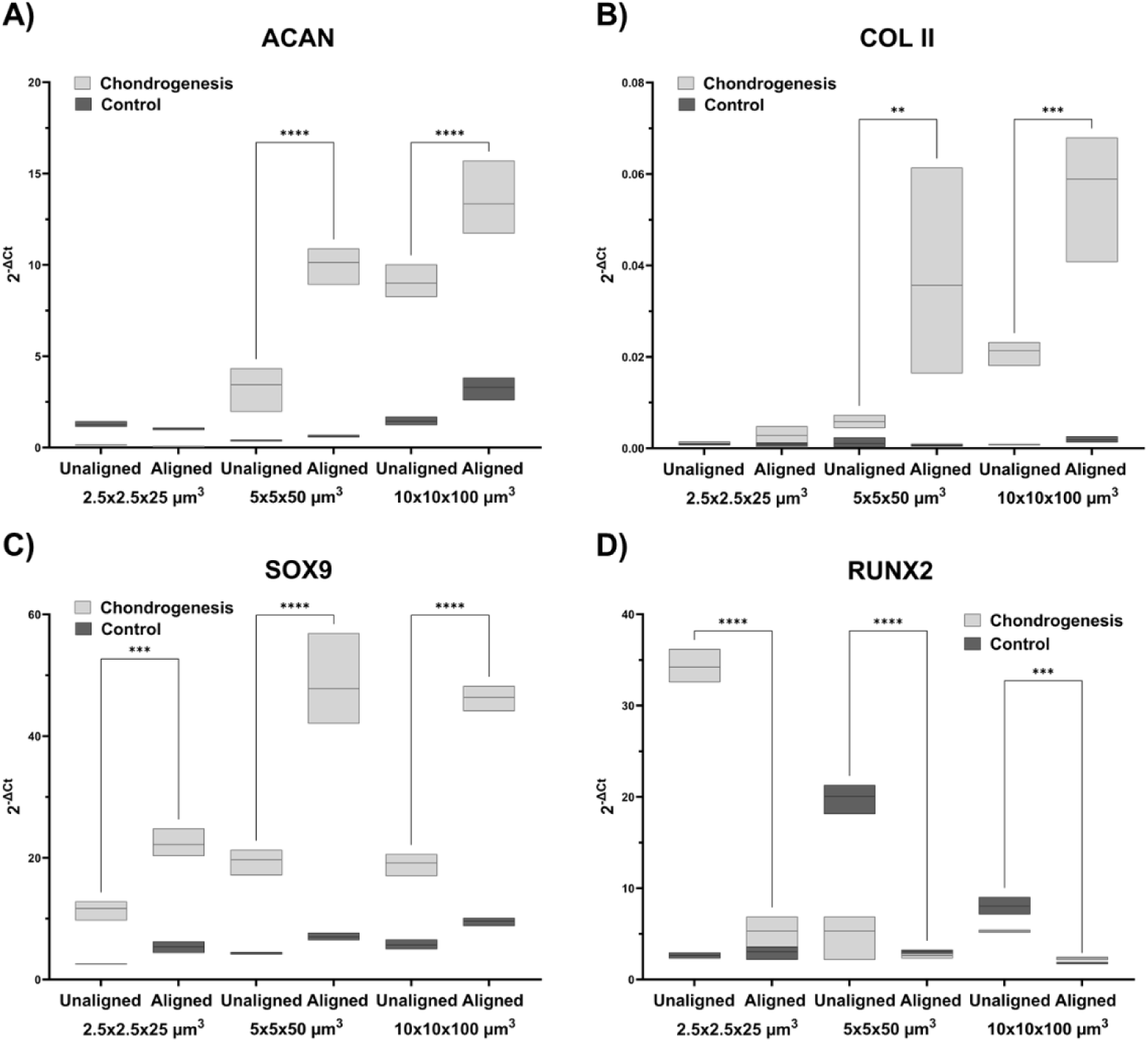
Chondrogenic and hypertrophic marker expression for different microgel dimension aligned vs. unaligned in control or chondrogenic conditions. Relative expression of A) *ACAN*, B) *COL2A1*, C) *SOX9*, and D) *RUNX2*. Data are presented as floating bars with a line indicating the mean. Samples come from n=3 technical replicates, from a n=10 pooled batch of hydrogels. A two-way ANOVA was used to evaluate the effects of differentiation and alignment on marker expression. Tukey’s post hoc test was applied for multiple comparisons. Significance is indicated as: ns (*p* ≥ 0.05), * (*p* < 0.05), ** (*p* < 0.01), *** (*p* < 0.001).

Additional to the previous chondrogenic markers, *RUNX2* was also evaluated as a marker for hypertrophic chondrocytes.^[47]^ *RUNX2* was upregulated in cells encapsulated with unaligned microgels cultured with chondrogenic differentiation media (**Figure 4 D**), indicating the biochemical stimuli provided by this media did not limit cells from hypertrophic chondrogenic differentiation.^[48]^ The alignment of the microgels led to a reduction of *RUNX2* expression, suggesting a prevention of chondrocyte hypertrophy and subsequent progression to endochondral ossification.^[49]^

Overall, the aligned microgels with the highest tested dimensions (10×10×100 µm^3^) showed an upregulation of chondrogenic markers (*SOX9*, *ACAN*, *COL2A1*) and reduced expression of the hypertrophic marker (*RUNX2*) compared to control hydrogels without microgels, unaligned microgels and smaller aligned microgel sizes (2.5×2.5×25 µm^3^, 5×5×50 µm^3^). The effect was independent of the donor of hMSCs, as similar trends were shown with two additional donors (see **Figure S2**). The dependency of chondrogenic differentiation on the size of the microgels could be attributed to the contributions of the stiff surface of the microgels. The employed microgels used have a previously reported elastic modulus of ∼400 kDa^[36]^, which is significantly higher that the surrounding hydrogel. Previous studies have shown that the addition of stiffening components (e.g., poly-l-lactic acid (PLLA) porous microspheres) to fibrous scaffolds led to increased new cartilage formation *in vivo* after 16 weeks.^[50]^ Additionally, anisotropic materials or composites with stiff elongated fiber-like structures have shown to guide hMSCs chondrogenic differentiation.^[51, 52]^ Our research indicates that the stiffening structures must have a minimum size to enhance chondrogenesis.

### 2.3 Osteochondral tissue repair with microgel in hydrogel system *in vivo*

The semi-orthotopic mouse model is a research tool to investigate the specific tissue niche at the cartilage and bone interphase and can be used to evaluate new methods to repair cartilage or osteochondral defects.^[38, 53]^ Briefly, osteochondral defects were created in biopsies of articular cartilage and underlying bone from bovine joints, after which the defects were filled with different hydrogel conditions and implanted subcutaneously in the back of NMRI-Fox1nu mice (**Figure 5 A**). The mice provide an *in vivo* environment to keep the tissue of the osteochondral biopsies alive, which enabled us to evaluate whether the Anisogel has the capacity to support cell infiltration and regenerate the damaged tissue. The different hydrogel and Anisogel conditions that were tested are specified in **Table 1**.

**Figure 5.**
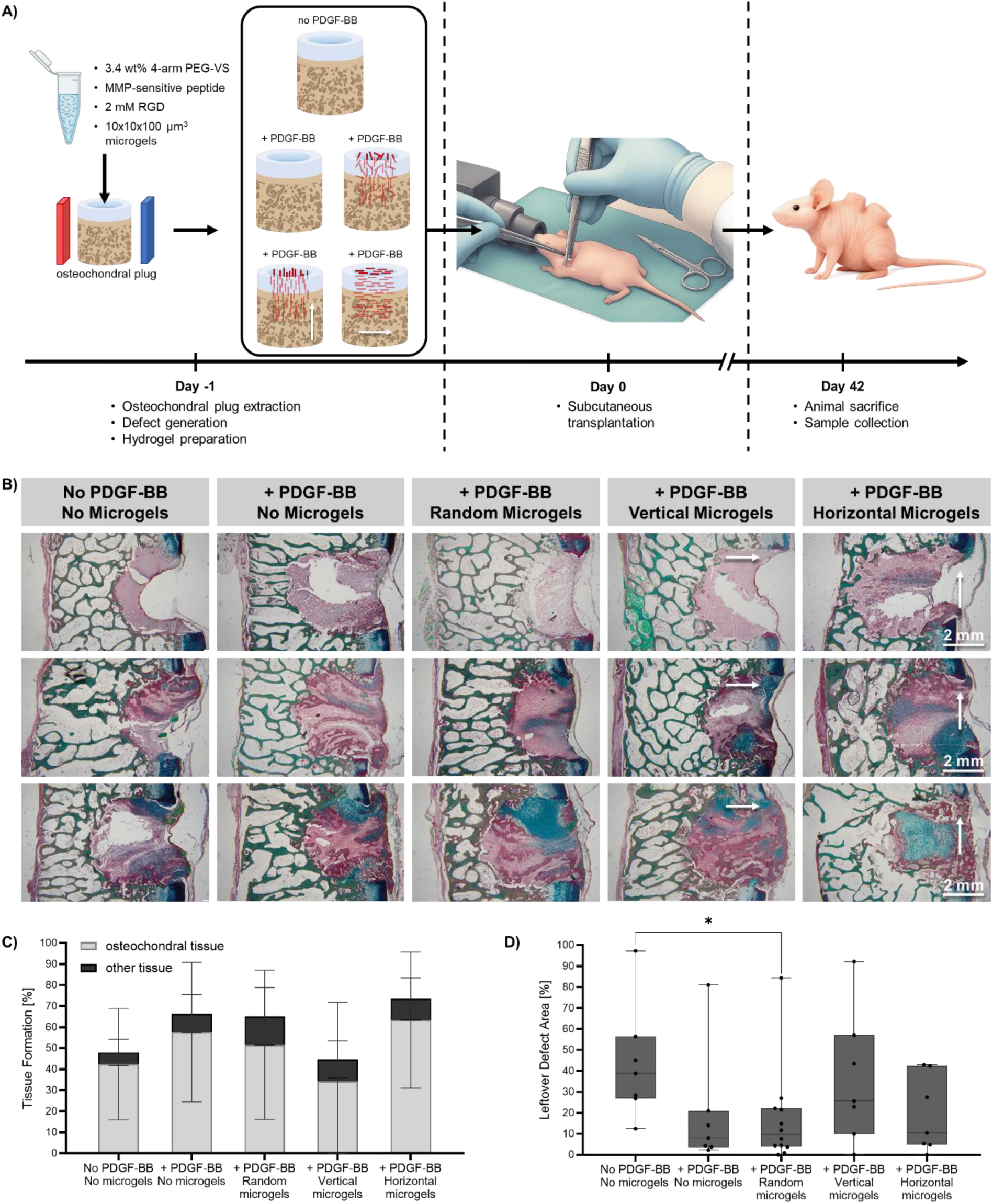
Optimized microgel-in-hydrogel system in semi-orthotopic mouse model *in vivo*. A) Schematic illustration of experimental conditions, timeline, and protocol. B) Three representative RGB stained sections of the biopsies per condition of osteochondral plugs retrieved 6 weeks post-implantation. Cartilage-like tissue is stained by Alcian Blue due to glycosaminoglycans (blue) and bone-like tissue stained by Picrosirius Red due to the mineralized matrix (deep purple). The arrows indicate the magnetic field direction. C) Bar plots display mean and SD for the quantification of the tissue formation in the defect space, divided into osteochondral tissue (gray) and other tissue (black). D) box plot showing the 1^st^ and 3^rd^ quartile, with bars depicting the maximum and minimum of the quantification of leftover defect area (not cell infiltrated gel and empty space). A Kruskal-Wallis test was used for evaluating significance is indicated as: * (*p* < 0.05).

**Table 1.** Overview of control and experimental groups for testing the Anisogel in semi-orthotopic mouse model.

|  | PDGF-BB loaded | Microgels included | Microgel orientation |
| --- | --- | --- | --- |
| Control group 1 | No | No | - |
| Control group 2 | Yes | No | - |
| Control group 3 | Yes | Yes | Random |
| Experimental group 1 | Yes | Yes | Horizontal → |
| Experimental group 2 | Yes | Yes | Vertical ↑ |

To promote endogenous cell migration, platelet-derived growth factor B (PDGF-BB) was added to the hydrogel precursor solution before gelation. PDGF-BB is known to enhance cell migration and proliferation.^[53, 54]^ Hence, it has been studied to improve *in vivo* cell infiltration in various hydrogels, such as PEG^[55]^, alginate^[56]^, fibrin^[57]^, or dextran hydrogels^[58]^. After 6 weeks, the samples were retrieved for histology and stained with hematoxylin-eosin (H&E, **Figure S3**) and Picrosirius Red-Fast Green-Alcian Blue (RGB) trichome staining (**Figure 5 B**).

In most samples, even the condition without PDGF-BB mixed in the hydrogel, the defect was filled with repair tissue after 6 weeks and no signs of inflammation were present (Figure S3 and Figure 5 B). This suggests that the modified PEG gel has a good biocompatibility. Small traces of left-over gel were visible in some defects, the majority of the gel was degraded and replaced by tissue within 6 weeks. While a lower average level of tissue formation was observed in the hydrogels without PDGF-BB, the addition of PDGF-BB in the hydrogel did not significantly increase tissue repair, possibly because of a very fast release of the growth factors from the gel (**Figure 5C**). We did observe a significantly higher left-over defect area without PDFG-BB, represented by regions of gel without cells and empty area, compared to the growth factor-loaded hydrogels with random microgels (**Figure 5 D**). The empty white area may appear due to hydrogel shrinking or tissue processing. This suggests that the PDGF-BB was stimulating the cells to infiltrate into the hydrogel, stabilizing the hydrogel during post-survival processing. Inclusion of microgels did not hamper tissue repair, albeit the vertical alignment seems to have a slightly lower defect filling with newly formed tissue (**Figure 5C**). This is likely because the defect size has the largest tissue interfacial surface at the sides of the defect and not at the bottom and the dense vertical oriented microgels might have blocked cell ingrowth from those sides. The studies were limited to 6 weeks, which is a relative short time to study repair with mature subchondral bone and articular cartilage.

Despite the initial promising results shown in the *in vivo* experiment, further optimization of the hydrogel is required to prove the potential of tissue guiding and differentiating structures, such as aligned microgels, in the osteochondral niche. Ongoing work is focused on integrated drug delivery systems to have a more sustained release of the growth factors. Moreover, the semi-orthotopic osteochondral defect model in mice is very useful to get a first impression of the capacity of a method to repair osteochondral defect tissue but lacks the synovial environment and mechanical loading and is prone to bone formation. To evaluate the full potency of selected conditions for cartilage regeneration, advanced translational studies using *ex vivo* bioreactors with mechanical loading, larger animal models, and better growth factor delivery approaches into our Anisogel system would be required.

## 3. Conclusion

In this study, we developed a microgel-in-hydrogel system to support anisotropic chondrogenic differentiation of human mesenchymal stem cells (hMSCs) *in vitro*, with the aim to induce cartilage formation *in vivo*. Through the process of biomaterial composition optimization, we identified that a 3.4 wt% MMP-degradable PEG-VS hydrogel modified with 2 mM RGD provided the most favorable conditions for cell ingrowth and chondrogenesis. The incorporation of anisotropy into the system, achieved by aligning magneto-responsive rod-shaped microgels of varying sizes, resulted in a significant and donor-independent enhancement of chondrogenic markers at the gene level (*SOX9*, *ACAN*, *COL2A1*), thereby highlighting the beneficial effect of anisotropic structural guidance on hMSC differentiation. Moreover, in a semi-orthotopic mouse model, encouraging preliminary findings demonstrated that cell ingrowth and tissue infiltration of osteochondral tissue in the defect area is not inhibited by the use of microgels, with only a slight non-significant decrease in the case of vertically aligned microgels. The delivery of PDGF-BB may have stimulated cell ingrowth but more sustained drug delivery systems are required. Therefore, to further support tissue repair, additional biochemical and biophysical cues must be considered, such as triggered growth factor release over a prolonged period of time. Moreover, future studies should focus on the generation of anisotropic hydrogel, with horizontal microgels in the surface layer, random microgels in the middle layer and vertical microgels in the deep layer, to better mimic articular cartilage with final testing in a larger animal model.

## 4. Experimental Section/Methods

### Cell culture of human mesenchymal stromal cells (hMSCs)

Bone marrow derived MSCs were purchased at passage 2 (19 year old male; additional experiments conducted with cells from 20 year old female / 25 year old male) from RoosterBio (Frederick, MD, USA) and sub-cultured in RoosterNourish™-MSC medium (RoosterBio) with 1 vol.% Penicillin/Streptomycin under a humidified atmosphere of 5% CO_2_ at 37 °C. For the following experiments, MSCs were used in passage 5. Before the cell encapsulation in hydrogels, the cells were detached with TrypLE™ Express (Gibco; Thermo Fisher Scientific) and resuspended in low-glucose DMEM (Dulbecco’s Modified Eagle Medium; Corning). After centrifuging at 300 rcf for 5 min, cells were concentrated to 2×10^6^ cells mL^-1^ in the final gel.

### Hydrogel formation

The hydrogel matrix components were diluted in low-glucose DMEM (Corning, pH=8) and mixed in a protein LoBind^®^ tube. Example values of the hydrogel mixtures can be found in the **Table S1** and **Table S2**. Briefly, polyethylene glycol vinyl sulfone (4-arm 20k, Creative PEGworks), the cell binding peptide (G**RGD**SPC, CPC Scientific) and the hMSCs suspension were mixed together. The MMP sensitive peptide (GCRE**GPQG`↓`IWGQ**ERC, GenScript) solution was prepared freshly and added immediately before casting. The gel precursor solution was mixed extensively and then pipetted to 15 μL droplets in a 48 well plate, which was flipped every 2 minutes until gelation to avoid cell sedimentation. After 45 min, 300 µL RoosterNourish™-MSC medium (RoosterBio) is added in every well and incubated at 37 °C and 5 vol% CO_2_.

### Microgel production

The microgels used for the Anisogel are produced by an in mold polymerization method, adapted from the technique previously reported by Babu *et al.*^36^ A silicon wafer patterned with the desired microgel dimensions was fabricated via photolithography (AMO GmbH) and subsequently treated with fluorosilazane. Polydimethylsiloxane (PDMS; Sylgard 184, Mavom) was mixed with its curing agent at a 10:1 ratio, poured over the wafer, and cured at 110 °C for 2 hours to create a mold. The polymer precursor solution was evenly spread across the PDMS mold using a polyethylene terephthalate (PET) foil (Goodfellow) placed on top of the liquid layer. The foil was then carefully peeled off to remove excess solution while avoiding bubble formation. The mold was subsequently exposed to UV light for 60 minutes in nitrogen atmosphere to initiate polymer cross-linking. The cured mold was coated with a layer of 12.5% (w/v) polyvinylpyrrolidone (PVP; 360 kDa, Sigma–Aldrich) dissolved in 99.9% isopropanol. After the PVP layer had fully dried, it was carefully peeled off, detaching the embedded microgels from the mold. The microgels were then released by dissolving the PVP in 99.9% isopropanol. The resulting suspension was washed three times with fresh 99.9% isopropanol, followed by sterilization in 70% isopropanol. Under aseptic conditions, the microgels were subsequently washed three times with Milli-Q water, once with sterile DMEM (Gibco), and then stored in DMEM supplemented 1% Pen/Strep (Gibco) at 4 °C until use. Microgels were counted using a Neubauer chamber and concentrated to 1 vol% in the final gel according to Rose *et al*..^44^ For further information on the microgel volume and size, please refer to **Table S3**.

### Anisogel formation

The hydrogel components were mixed as described in the previous section. The right concentration of microgels (see **Table S4**) was mixed prior to casting. To achieve microgel alignment, the well plate was placed on a well plate formed, self-designed magnet, which is kept on the well plate during the gelation time. The Anisogels were also flipped every 2 min to avoid microgel and cell sedimentation. After 45 min, the aligned microgels were removed from the magnet plate and 300 µL RoosterNourish™-MSC medium (RoosterBio) was added in every well and incubated at 37 °C and 5 vol% CO_2_.

### Cell culture in PEG-VS hydrogels and Anisogels

From the first day on after gelation, chondrogenesis is started for 21 days with media being changed three times per week. The control medium consists of low-glucose DMEM (Corning), 10 vol% heat-inactivated FBS (Gibco; Thermo Fisher Scientific), 1 vol% Penicillin/Streptomycin, and 1 vol% GlutaMax 100X (2 mM; Gibco; Thermo Fisher Scientific). The basal media for chondrogenesis consists of high-glucose DMEM (Gibco; Thermo Fisher Scientific), 2 mM GlutaMax 100X (Gibco; Thermo Fisher Scientific), 1 vol% Penicillin/ Streptomycin (Pen/Strep; Gibco; Thermo Fisher Scientific), 2.5·10^-4^ M Ascorbic acid (Honeywell Fluka), 40 µg mL^-1^ L-proline (Thermoscientific), 1 mM Sodium pyruvate (Gibco; Thermo Fisher Scientific), and 1 vol% Insulin-transferrin-selenium (ITS; Gibco; Thermo Fisher Scientific). The chondrogenic medium is prepared freshly every time before medium change. Therefore, 10 ng/mL TGF-β1 (AcroBiosystems) and 10^-7^ M dexamethasone (Thermoscientific) is added to the basal medium.

### Immunofluorescent Staining and Imaging

After the 21 days of encapsulation, cells in the hydrogels were stained for further confocal imaging. Initially, the hydrogels were washed with PBS and then fixed for 40 min with 4% PFA (Sigma Aldrich). After washing with PBS, the samples were permeabilized with 0.1% Triton X-100 (in PBS, Sigma Aldrich) for 20 min at RT. Afterwards, the samples were incubated in a blocking solution of 4% BSA and 0.1% Triton X-100. Then, the samples were incubated overnight at 4 °C with the primary antibodies: collagen II (0.5µg/mL, mouse IgG2a, MA1-40065, Invitrogen) and aggrecan (2,75µg/mL, rabbit IgG, 30532-1-AP Proteintech). After the incubation with the primary antibodies, the samples were washed with PBS, two times at RT to remove the unreacted primary antibody. The staining of phalloidin (1:1000, AAT Bioquest), and secondary antibody: AF647 anti rabbit (1:200, Invitrogen) or AF 594 anti-mouse (1:200, Invitrogen) was carried out overnight at 4 °C. Prior to imaging, samples were stained with DAPI (1:200, Sigma Aldrich) and washed with PBS. Imaging was conducted at the Opera Phenix™ Content Screening System.

### Image Analysis

Superimposed z-stack images were analyzed using Fiji (version 2.17.0)^[59]^ by converting the images to 8-bit, setting the threshold to convert the images into black and white images, and applying the watershed function (for separation of connected cells). Nuclei were counted, and protein area was measured by using the Analyze Particle function. The obtained area values were then normalized by dividing by the corresponding nuclei number.

For alignment analysis, a custom-made script was used based on previous work.^[28]^ The alignment of the microgels was quantified using an orientation analysis on the images using an elliptical Mexican hat filter.^[60]^ In summary, the procedure was as follows: Images were loaded and background corrected. Background images were created convolving the image with a Gaussian kernel with a width (standard deviation) of 10 pixels, window size of 61 pixels. This background was subtracted, and all negative values were set to zero. Then the image was smoothed with another Gaussian kernel (width of 0.5 pixels, window size of 11 pixels) again applying convolution. Because the images had relatively broad structures, an edge detector filter was then used, based on the first derivative of a Gaussian kernel. Orientation was detected using an elliptical Mexican hat filter, which is the negative of a second derivative Gaussian with a width of 10 pixels in one direction and 1 pixel perpendicular to it, and a window size of 41 pixels. The kernels are normalized to have a sum of 1. This ellipsoid ‘hat’ kernel was rotated to 20 angles in the [0, 180) degrees range and a maximum intensity was recorded with the corresponding angle for every pixel. Then a threshold was generated to distinguish objects from background using Otsu’s method, and the detected object pixels and their corresponding orientation values stored. Histograms were created counting the number of pixels per rotation angle. The full width of half maximum was determined using a linear interpolation of the two neighboring points around the half of the maximum count.

### Rheology

Rheological measurements were carried out using a Discovery HR-3 hybrid rheometer equipped with a 20 mm cone-plate geometry (2°). For each experiment, 74 μL of the hydrogel mixture was pipetted onto the rheometer plate maintained at 37 °C. Time-dependent measurements were recorded over 60 minutes at a frequency of 0.5 Hz, with an oscillatory strain of 2% and a gap size of 50 μm, performed in triplicate for each condition.

### Quantitative Polymerase Chain Reaction (qPCR)

Before the RNA extraction, the encapsulated hMSCs were extracted by washing the hydrogels once with PBS, followed by a 10 min incubation in TrypLE™ Express (Gibco; Thermo Fisher Scientific) to digest the surrounding hydrogel. The suspension was collected into falcons, resuspended using a syringe and cannula (0.8 x 120 mm) to mechanically disrupt the matrix, and centrifuged at 400 rpm for 5 min at RT. The supernatant was discarded and the pellet was resuspended in 700 µL of lysis buffer (79216, Qiagen) and left on ice for 10 min. The lysates were centrifuged at 16.000 g, 4 °C for 4 minutes. The supernatant was transferred to a cell-lysate homogenizer (QiaShredder, Qiagen) and used according to the manufactureŕs instructions. The flow through was collected and used for RNA extraction.

Total RNA was extracted from the cell lysates using the RNeasy Mini Kit (Qiagen) following the manufacturer’s protocol. RNA concentration and purity were assessed using a NanoDrop spectrophotometer (Thermo Fisher Scientific). For each sample, 500 ng to 1 µg of total RNA was reverse transcribed into cDNA using the PrimeScript™ RT Master Mix (RR036A, TaKaRa) according to the manufacturer’s instructions.

Quantitative PCR was performed using (CFX384 Real-Time Systems, Bio-Rad) with SYBR Green-based detection (1725270, Bio-Rad). Each 10 µL reaction contained 8 µL of SYBR Green Master Mix (Bio-Rad), 0.5 µM each of forward and reverse primers (see **Table 2**), and 2 µL of diluted cDNA template. The thermal cycling conditions were as follows: initial denaturation at 95 °C for 3 minutes, followed by 40 cycles of denaturation at 95 °C for 10 seconds, annealing in a gradient temperature from 57-65 °C for 30 seconds, and extension at 72 °C for 30 seconds, with a melt curve analysis performed at the end to confirm specificity of amplification.

**Table 2.**
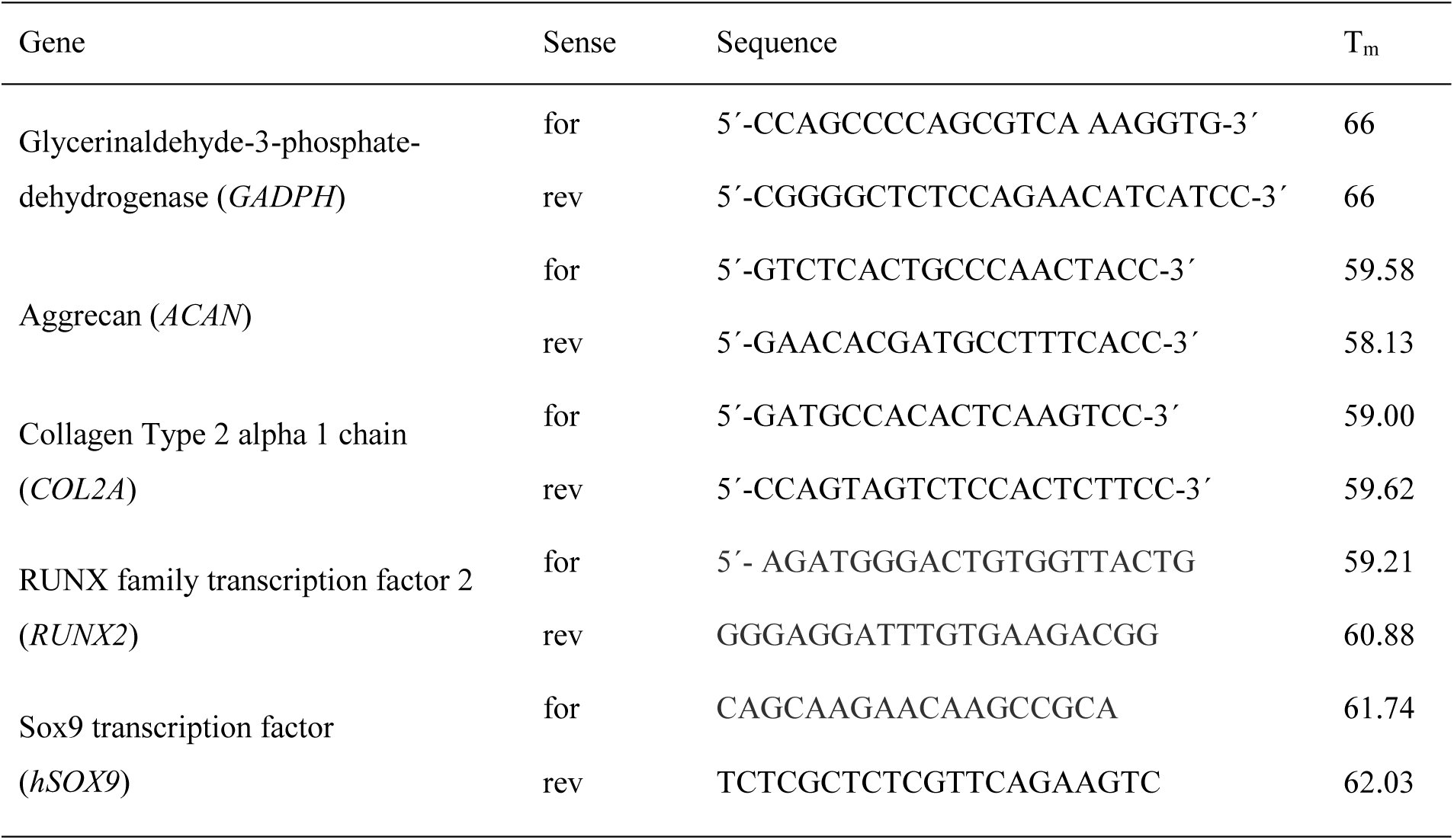
Sequences and melting temperatures of used forward and reverse primers.

All reactions were run in triplicate (technical replicates), and DNA-free water controls were included to rule out contamination. The gene expression levels were normalized to the expression of the housekeeping gene, *GAPDH*, which was constant among all conditions and experimental groups. The values were reported as relative quantification was calculated using the 2^−ΔCt^ method.

### In vivo experiments

To assess the impact of optimized microgel-in hydrogel system on cartilage repair in an osteochondral tissue environment, the established semi-orthotopic mouse model was applied.^[54]^ Fresh osteochondral biopsies (8 mm diameter, 5 mm height) were harvested from bovine metacarpal-phalangeal joints (LifeTec, Eindhoven) and osteochondral defects were created on the cartilage side of the biopsies (4 mm diameter, 4 mm height). These defects were filled with the different biomaterials according to the five conditions tested (**Table 1**).

The defects in the osteochondral biopsies were filled up with the hydrogel precursor solution, and the certain microgel alignments were achieved by different magnetic fields. After one-hour gelation time, these biopsies were cultured in high glucose DMEM (4.5 g/L glucose, Gibco) supplemented with 10% fetal bovine serum (FBS, Gibco), 50 μg/mL gentamycin (Gibco), and 1.5 μg/mL fungizone (Gibco) at 37 ℃ overnight.

This animal experiment was approved by the Ethics Committee for Laboratory Animal Use (Study Nr: 2115392-01). Ten 10-week-old NMRI-Fox1nu female mice (Janvier Labs, St. Berthevin, France) were randomly assigned and housed under specific pathogen-free conditions with a regular day/night light cycle and allowed to adapt to the conditions of the animal facility for 7 days. Food and water were available ad libitum. One day after the gelation, the cartilage side of the osteochondral biopsies was covered with a circular 8 mm Neuro-Patch membrane (Braun) to prevent the ingrowth of host cells from the top. To create space for the implant, four small incisions are made on the back of the mice. A subcutaneous pocket is then created using blunt dissection. The biopsies were randomly implanted in pockets under 2.5–3% isoflurane anesthesia (1000 mg/g body weight, Laboratorios). One osteochondral biopsy was implanted per pocket, and four osteochondral biopsies were implanted per mouse. The wounds were closed with two staples per wound (Autoclip).

At 1 h before surgery and at 6–8 h after surgery, 0.05 mg/kg body weight of buprenorphine (Temgesic, Reckitt Bensicker) was injected subcutaneously to ensure pre- and postoperative analgesia. After emerging from anesthesia, the mice were closely monitored for stress and other discomfort and weight was checked periodically. Mice were always housed in the same group (3-4 mice per cage). One mouse was excluded from the study after 3 weeks due to the impaired wound healing, and was euthanized by cervical dislocation under 2.5-3% isoflurane anesthesia. After 6 weeks, the remaining ten mice were euthanized using the same method, and the osteochondral biopsies were harvested. All samples were fixed in 4% formalin for further analysis. Ultimately, seven biopsies per group were obtained from control groups 1 and 2 and experimental groups 1 and 2, while twelve biopsies were collected from control group 3.

### Histology staining

All samples were decalcified with 10% EDTA, embedded in paraffin and sectioned at 6 μm. Three sections localized in the shallow, middle and deep layers (each separated by 450 μm) within the defect area of each sample were collected for histology staining. The cartilage and bone tissue formation was assessed by histology staining using H&E staining and RGB trichromestaining.^[61]^ The classification and quantification of different categories in RGB staining was performed using QuPath software (v0.6.0). Briefly, the areas of newly formed cartilage tissue, bone tissue, other tissue, gel with infiltrated cells without tissue formation, and leftover (sum of gel without cells and empty area) were manually classified (**Figure S4**, **Figure S5**) and their areas calculated separately. The percentages were calculated as the ratio of these categories to the total defect area. All slides were quantified by one investigator blinded to the experimental conditions.

## 5. Acknowledgements

We acknowledge the funding from Werner Siemens Foundation (WSS) for the project TriggerInk.

## Data Availability Statement

The data that support the findings of this study are available from the corresponding author upon reasonable request.

## Supporting Information

**Table S1.** Example values for the production of 300 µL the hydrogels with variations in RGD concentration.

| | 150 $\mu\text{M}$ | 500 $\mu\text{M}$ | 1 mM | 2mM |
| --- | --- | --- | --- | --- |
| Cells | 118.6 $\mu\text{L}$ | 112.0 $\mu\text{L}$ | 102.5 $\mu\text{L}$ | 83.5 $\mu\text{L}$ |
| PEG-VS 4arm, 20k, 20% | 84.4 $\mu\text{L}$ | 87.2 $\mu\text{L}$ | 91.1 $\mu\text{L}$ | 99.0 $\mu\text{L}$ |
| GRGDSPC (25 mg/mL) | 1.7 $\mu\text{L}$ | 5.5 $\mu\text{L}$ | 11.1 $\mu\text{L}$ | 22.2 $\mu\text{L}$ |
| MMP sensitive peptide (3 w/v%) | 95.3 $\mu\text{L}$ | 95.3 $\mu\text{L}$ | 95.3 $\mu\text{L}$ | 95.3 $\mu\text{L}$ |

**Table S2.** Example values for the production of 300 µL of hydrogel with variations in the PEG-VS concentration.

|  | 3.4 wt% | 5 wt% | 6.5 wt% | 10 wt% |
| --- | --- | --- | --- | --- |
| Cells | 172 $\mu\text{L}$ | 130.7 $\mu\text{L}$ | 90.7 $\mu\text{L}$ | 65.9 $\mu\text{L}$ |
| PEG-VS 4arm, 20k, 20% | 59.4 $\mu\text{L}$ | 81.6 $\mu\text{L}$ | 101.1 $\mu\text{L}$ | 146.4 $\mu\text{L}$ |
| GRGDSPC (25 mg/mL) | 19.2 $\mu\text{L}$ | 19.2 $\mu\text{L}$ | 19.2 $\mu\text{L}$ | 19.2 $\mu\text{L}$ |
| MMP sensitive peptide | 49.4 $\mu\text{L}$<br>(3%) | 68.5 $\mu\text{L}$<br>(3%) | 89.0 $\mu\text{L}$<br>(3%) | 68.5 $\mu\text{L}$<br>(6%) |

**Table S3.** Microgel concentrations in hydrogel for 1 vol% dependent of the microgel size.

| Microgel size | # of microgels per $\mu\text{L}$ hydrogel |
| --- | --- |
| 2.5x2.5x25 $\mu\text{m}^3$ | 64 000 #/$\mu\text{L}$ |
| 5x5x50 $\mu\text{m}^3$ | 8 000 #/$\mu\text{L}$ |
| 10x10x100 $\mu\text{m}^3$ | 1 000 #/$\mu\text{L}$ |

**Table S4.** Example values for the production of 300 µL of hydrogel with variations in the microgel size.

|  | 3.4 wt% |
| --- | --- |
| Microgels | 100 $\mu\text{L}$ |
| Cells | 72.0 $\mu\text{L}$ |
| PEG-VS 4arm, 20k, 20% | 59.4 $\mu\text{L}$ |
| GRGDSPC (25 mg/mL) | 19.2 $\mu\text{L}$ |
| MMP sensitive peptide 3% | 49.4 $\mu\text{L}$ |

**Figure S1.**
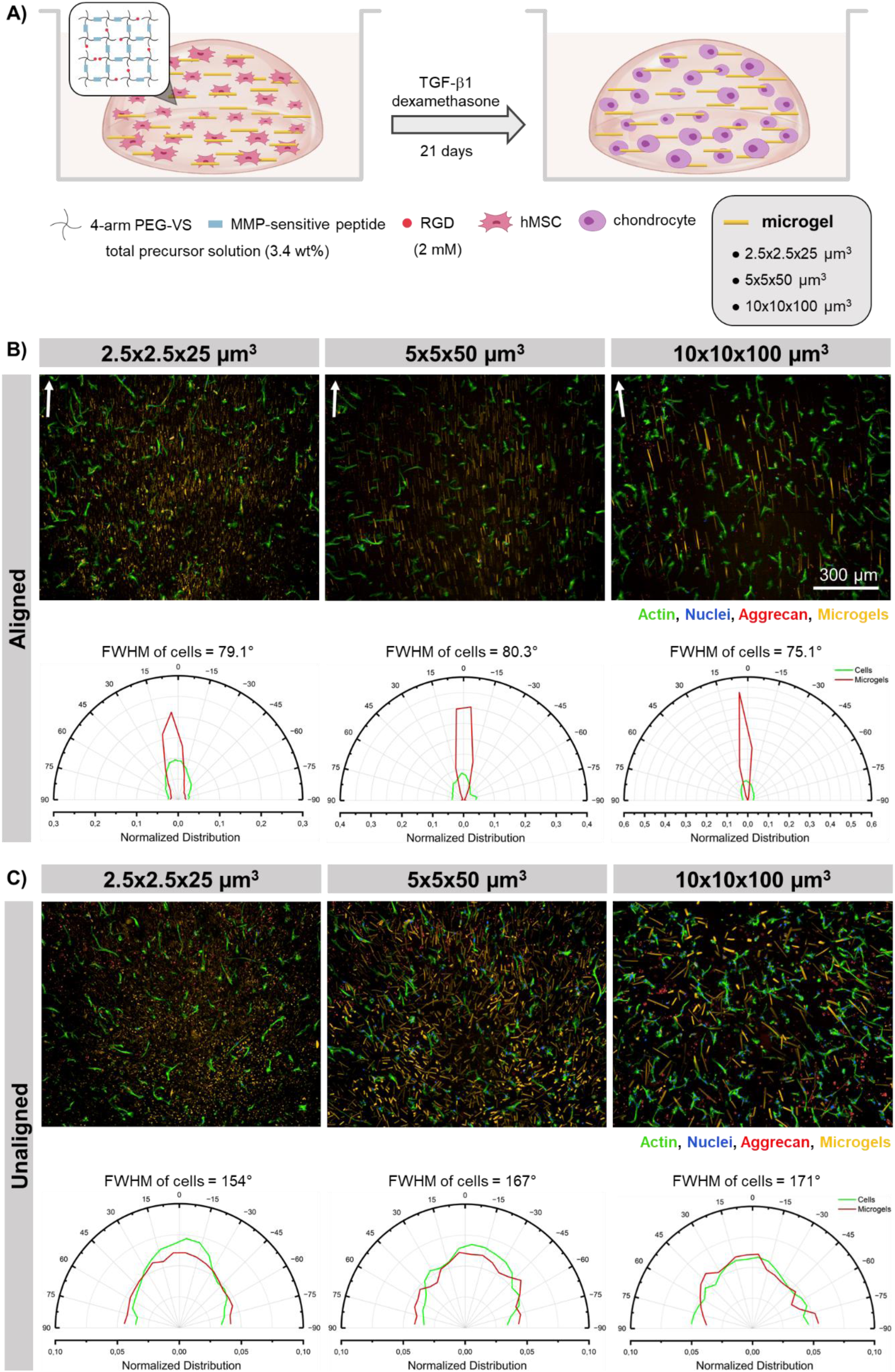
Varying the microgel dimensions and alignment of PRINT microgels in optimized PEG-VS hydrogel. A) Schematic protocol of hMSC differentiation in optimized PEG-VS hydrogel droplets for microgels with dimensions of 2.5×2.5×25 µm^3^, 5×5×50 µm^3^, or 10×10×100 µm^3^. Representative confocal microscopy images of hMSCs in hydrogels stained with F-actin (green) and DAPI (blue) cultured in optimized PEG-VS hydrogels in control media used for the correspondent angle histograms depicting the full width at half maximum (FWHM, of n=1 hydrogel containing n>80 cells in the field of view) of cell (green) and microgel (red), in aligned (B) and unaligned (C) microgels.

**Figure S2.**
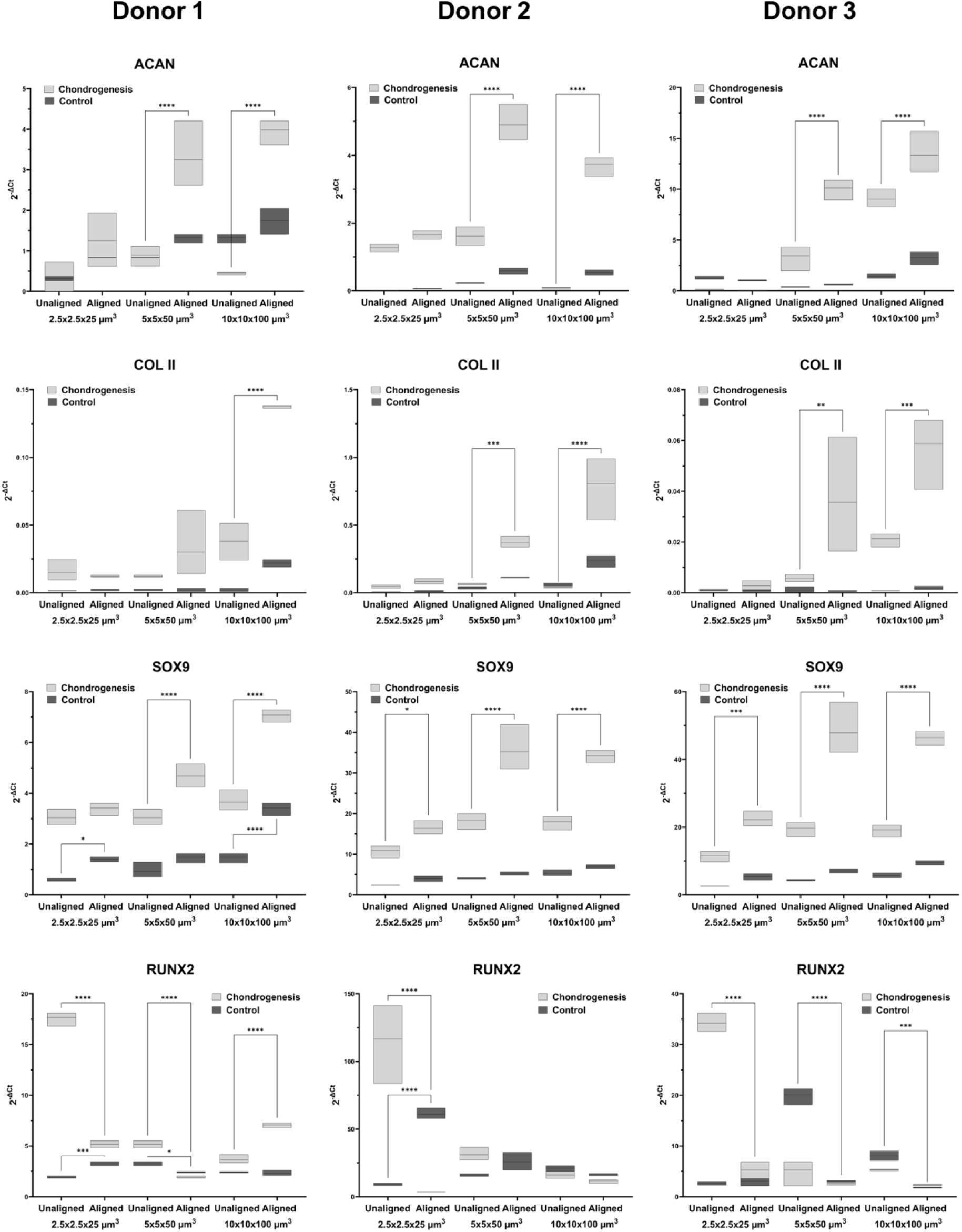
Chondrogenic and Osteogenic marker expression for different microgel dimension aligned vs. unaligned in control or chondrogenic conditions for 3 different hMSC donors. Donor 1 (20 year old female), Donor 2 (19 year old male) and donor 3 (25 year old male). Relative expression of A) Aggrecan, B) Collagen II, C) SOX9, and D) RUNX2. Data are presented as floating bars with a line indicating the mean. A two-way ANOVA was used to evaluate the effects of differentiation and alignment on marker expression. Tukey’s post hoc test was applied for multiple comparisons. Significance is indicated as: ns (*p* ≥ 0.05), * (*p* < 0.05), ** (*p* < 0.01), *** (*p* < 0.001).

**Figure S3.**
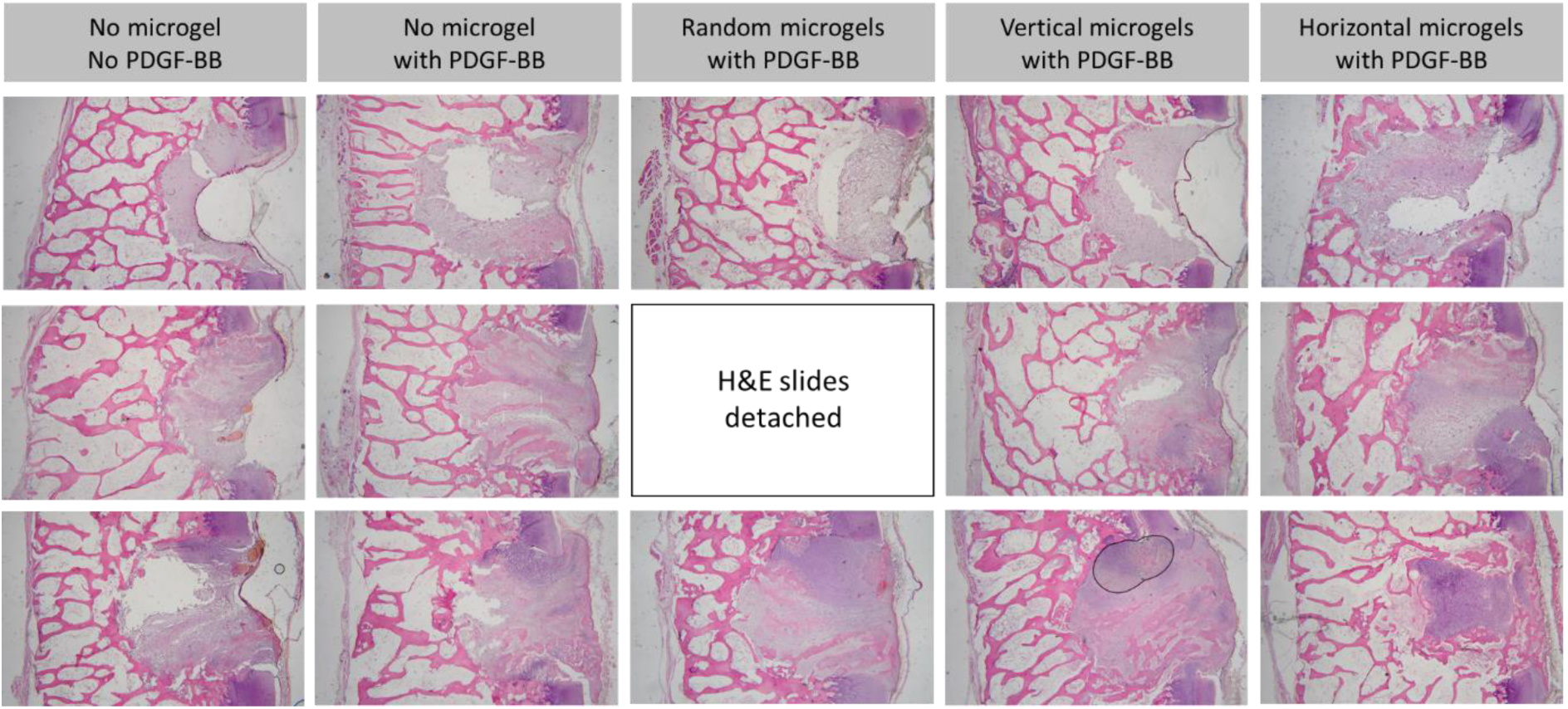
Hematoxylin-eosin staining of osteochondral plugs. Three representative images per condition of retrieved osteochondral plugs from a 6-week *in vivo* mouse orthotopic model representing the worst condition in the top, average in the middle and the best condition in the bottom row.

**Figure S4.**
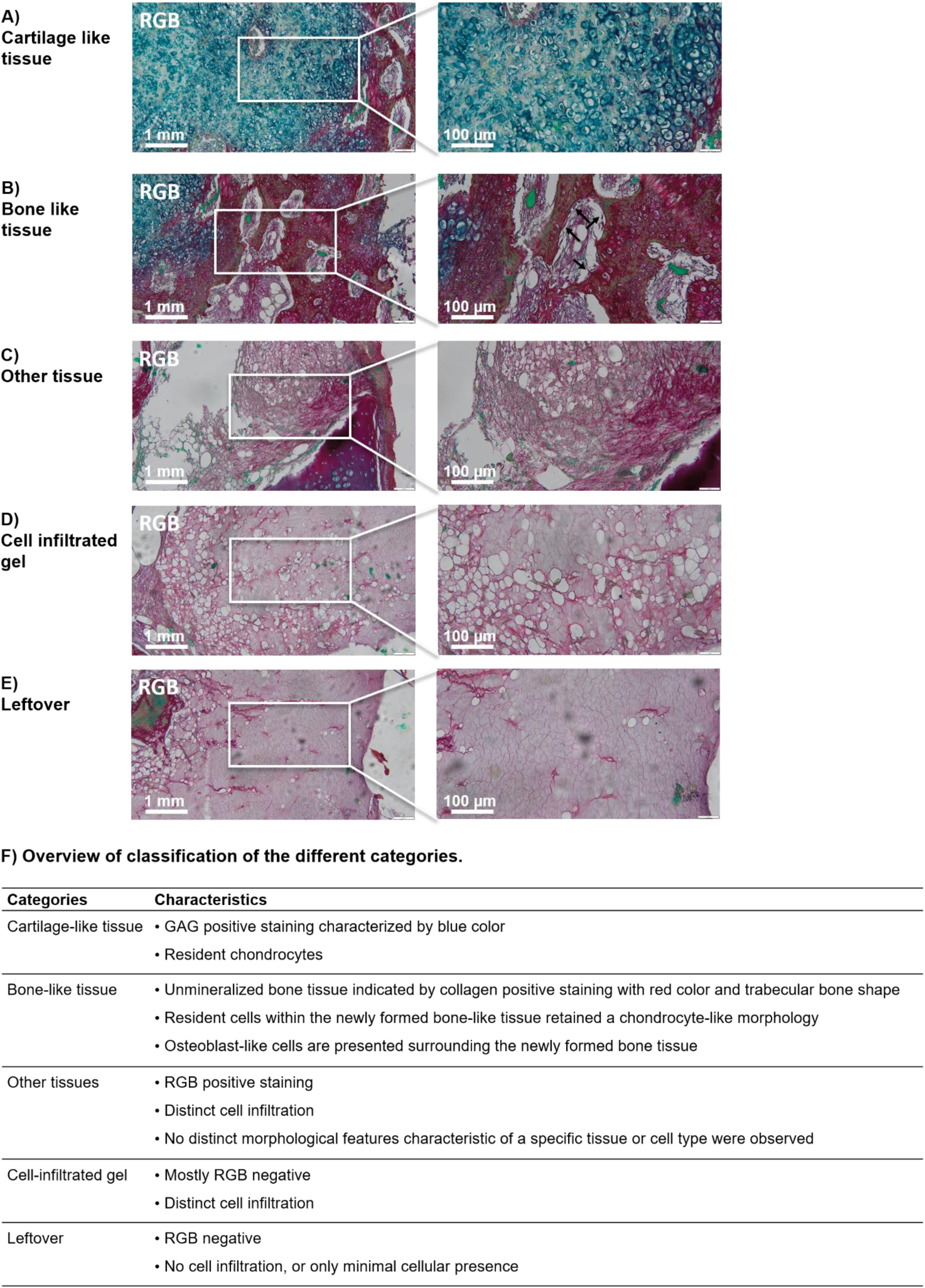
RGB trichrome staining of osteochondral plugs and tissue classification. Classification of the tissue/material in the defect area: A) Cartilage-like tissue stained by Alcian Blue due to glycosaminoglycans, B) bone-like tissue stained by Picrosirius Red due to the mineralized matrix, c) other tissues, sections with unclear morphological characteristics, d) cell infiltrated gel, hydrogel with single cells but not defined tissue formation and e) leftovers or cell free gel. F) Definition of classification in categories.

**Figure S5.**
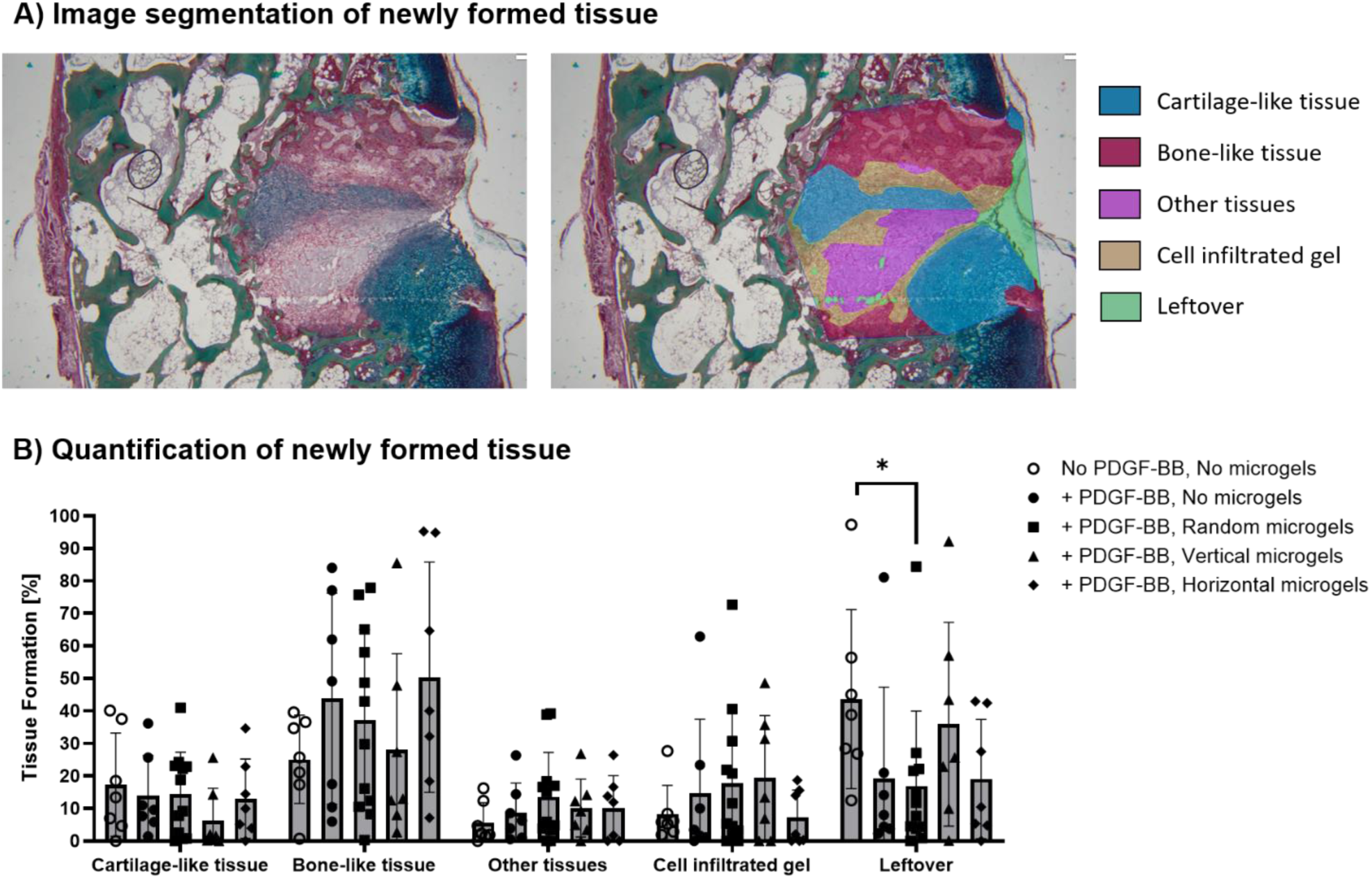
A) Example of an image segmentation of a plug with representative tissue types described in Figure S4. B) Individual quantification of the previously mentioned tissue types using the RGB staining.

